# Holographic optical trapping Raman micro-spectroscopy of interacting live cells

**DOI:** 10.1101/292334

**Authors:** F. Sinjab, H. M. Elsheikha, D. Awuah, G. Gibson, M. Padgett, A. M. Ghaemmaghami, I. Notingher

## Abstract

We present a combined holographic optical tweezers and confocal Raman spectroscopy instrument which allows fast, flexible, and interactive manipulation with molecular measurement of interacting live cell systems. Multiple laser foci created using a spatial light modulator are simultaneously used for optical trapping and spontaneous Raman spectroscopy. To enable confocal Raman measurements with high spectral resolution, a digital micro-mirror device was used to generate reflective pinholes which are matched to each laser trap. We demonstrate this unique capability by initiating complex interactions between multiple live cells whilst non-invasively acquiring Raman spectra of the processes with high spatial, spectral, and temporal resolution.

## INTRODUCTION

Molecular transport underpins all cell-cell communication and interactions in the life sciences. This involves complex spatially and temporally varying chemical processes responsible for many fundamental biological phenomena. Examples include the invasion of pathogenic species into host systems and the communication between cells involved in the innate and adaptive immune responses. Detailed measurement of such phenomena is important both for furthering basic understanding of biophysical processes and in practical applications such as engineering immuno-modulatory strategies for autoimmune diseases, cancers and vaccinations. Studying such complex systems requires the ability to manipulate cells without comprimising their viability or functional properties. At the same time, tools are needed which can non-invasively measure the underlying molecular properties of cells to detect changes in their biochemical characteristics over time.

Raman micro-spectroscopy (RMS) is a sensitive label-free analytical technique which is well suited to measuring biochemical signals with high chemical specificity and diffraction-limited spatial resolution (1,2). Within the scope of cell biology, RMS has been applied to the study of a wide range of species, from single bacteria (3,4), to fungal (5) and animal cells (6-8). RMS has also been applied to the study of cell systems relevant to medicine, including immunology (9,10), parasitology (11,12), stem cell biology (13,14), and cancer cell biology (15,16). RMS measurements usually involve tightly focusing a laser beam at a point on a sample, which excites spontaneous Raman scattering from the vibrational modes of the molecules in this region. A suitable technique for controlling live cells is optical trapping/tweezing(17,18), which has been demonstrated for a variety of applications across the life sciences (19,20). Modern holographic optical trapping (HOT) instruments are capable of generating many simultaneous trapping beams using liquid-crystal spatial light modulators (LC-SLM), which allow interactive control in real-time(21).

Optical trapping is a somewhat natural partner for RMS of single cells, as both techniques share many of the same requirements, such as the use of near-IR wavelengths to avoid absorption-induced damage, and improved performance with higher numerical aperture microscope objectives. Using a single focused laser beam, optical trapping has been combined with Raman spectroscopy for many biological applications, from the identification of bacterial spores to Raman activated cell sorting (4, 15, 16, 22-30). Single-beam optical trapping is compatible with the instrumentation used for RMS which typically involves a confocal pinhole and/or a narrow spectrometer entrance slit to facilitate high spectral resolution. However with the multi-point HOT approach, a single pinhole and/or slit is not suitable as the Raman scattering from most positions would be prevented from reaching the detector. Several approaches towards reconciling this problem have been proposed, such as using a rapidly scanned laser beam dwelling at multiple locations for time-shared multi-point measurement with a matched mirror to align sampling points into a spectrometer(29), power-shared excitation with an LC-SLM using a fixed array of electronically controlled pinholes(31) or imaging directly onto a detector for small highly scattering objects(32).

In order to fully utilize the flexibile laser-trap generation of the LC-SLM used for HOT in RMS, it is desirable to have a similarly rapid and flexible tool for producing an array of matched pinholes. One method from fluorescence microscopy involves the use of a programmable digital micro-mirror device (DMD), which functions as a binary-choice reflective-mode SLM by directing incident light into one of two directions. The use of a DMD has been demonstrated both for confocal(33) and wide-field(34) fluorescence microscopy. Using this approach, reflective pinholes can also be generated to collect the Raman scattering from sampling positions generated by an LC-SLM(35).

Here, we describe a video-rate HOT-RMS instrument with detection using reflective pinholes on a DMD. This instrument can manipulate and simultaneously measure Raman spectra from several optically trapped objects. We demonstrate the capability and limits of the instrument for measuring RMS spectra from moving trapped polymer beads, as well as various power considerations for measurements of multiple trapped cells. Finally we demonstrate two examples of multi-cellular interactions involving HOT control and RMS readout, first on the formation of an immunological synapse between T cells and dendritic cells (DCs), and second a host-pathogen interaction involving the parasite *Toxoplasma Gondii* and human brain microvascular endothelial cells (hBMECs).

## HOT-RMS INSTRUMENTATION

A schematic description of the HOT-RMS instrument is presented in Figure 1, the details of which are described in the supplementtary materials. A CW laser with power *P_0_* is shared into multiple laser foci each of power *P_i_* at the microscope sample plane by displaying a phase hologram at the LC-SLM using the RedTweezers software developed by Bowman et al.(21). Each laser focus can be controlled independently, simultaneously functioning as an optical trap and spontaneous Raman excitation source. Backscattered Raman photons are collected by the same 1.2 NA microscope objective. For HOT-RMS measurements, a filter wheel containing a NIR reflective mirror (M1 in Figure 1) directs the Raman light out of the microscope towards a Raman spectrometer. Before the Raman photons reach the spectrometer, they should ideally be spatially and spectrally filtered (the latter is carried out using a bandpass or notch filter to reject excitation wavelengths). Spatial filtering serves multiple purposes: first, any undesirable out-of-focus or stray light is rejected before entering the spectrometer (i.e. confocality is increased), and second, the spectral resolution is improved to resolve Raman linewidths with minimal spectral overlap. Typically this filtering is carried out using a spectrograph entrance slit (to ensure high spectral resolution) and/or a confocal pinhole, before being dispersed within the spectrometer.

**Figure 1:**
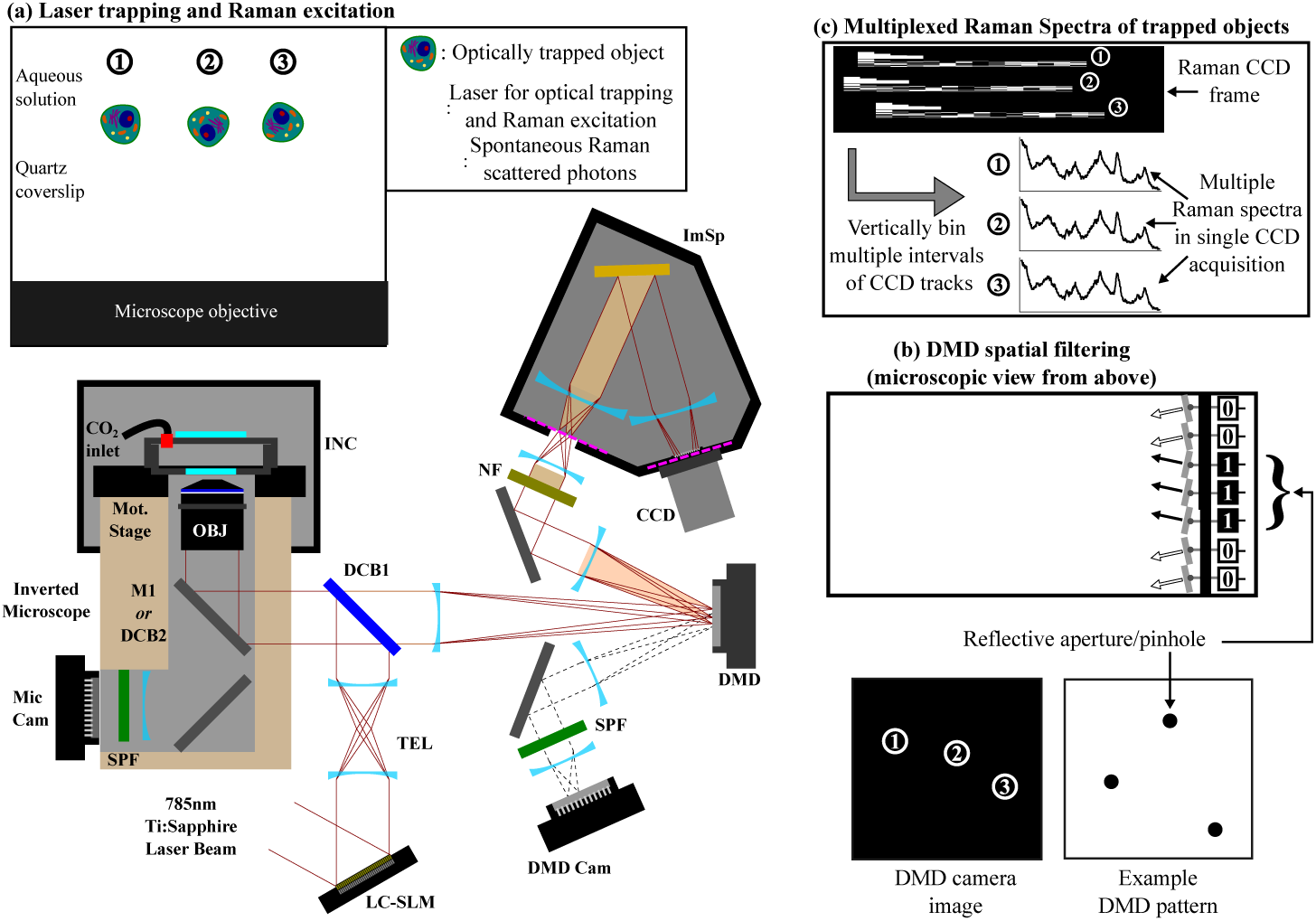
HOT-Raman instrument. (a) Multiple laser foci are used to trap and excite Raman scattering from micro-objects on a microscope. (b) Raman scattering is spatially filtered by a DMD. (c) Raman spectra from each trap can be read-out simultaneously. Glossary: **LC-SLM**: liquid-crystal spatial light modulator, **INC**: microscope incubator enclosure, **TEL**: 4f telescope relay, **DCB**: dichroic beamsplitter, **OBJ**: microscope objective, **DMD**: digital micro-mirror device, **ImSp**: imaging spectrometer, **DMD Cam**: CMOS inspection camera for DMD, **SPF**: short-pass filter, **NF**: notch filter, **M**: near infra-red mirror, **Mic Cam**: microscope camera. Dashed pink lines represent planes perpendicular to the optical axis which are sample-conjugate. Dashed black lines show rays reflected from the DMD plane away from the spectrometer direction.

Here a DMD is instead used for spatial filtering. This functions by creating a binary image of reflective apertures in software (written as a LabVIEW plugin for the RedTweezers control software) which can be calibrated to match 1-to-1 with each laser trap co-ordinate in the sample-conjugate plane at the DMD display. The DMD also has the advantage of choosing an aperture size in software, allowing simple optimization of the spectral resolution and confocalality vs. optical throughput (see Figure S1). The effect of spatial filtering also depends on the physical propeties of the sample itself. For example, degradation of spectral resolution is more apparent when measuring bulk samples compared to microscopic particles (see Figures S1 and S2). In order to match the DMD pinhole positions to the laser trap co-ordinates, polystyrene micro-particles were optically trapped across the usable field-of-view (FOV), and were monitored with the DMD camera and real-time Raman spectra. These readouts were used to assess the overlap between the DMD pinhole and image of the micro-particles, and the Raman signal strength respectively. Typical DMD-pinholes diameters of 110μm were found to be optimal for maintaining high spectral resolution and optical throughput. The axial position of a trapped object was also optimized using LC-SLM control (see Figure S3), and abberration correction of the laser traps was carried out using the RedTweezers software options (see Figure S4). Finally, to demonstrate dynamic HOT-RMS measurement capability, Raman spectra were rapidly acquired from four trapped polystyrene spheres simultaneously, which were manipulated interactively in real-time at the sample. This was achieved using four distinct regions of the CCD, corresponding to four sample-conjugate regions. Frames from each CCD region were binned in hardware allowing independent readout at 10 Hz (corresponding to 40 spectra per second), for a total time of 100s. During this time, the four trapped particles were manipulated around the four FOV regions, with microscope illumination from a halogen lamp for the first 90s. Figure 2 shows video frames from the experiment, with the corresponding simultaneous time-course Raman spectra. Figure 2 shows that the manipulation of a trapped particle within a particular FOV can result in a spectral shift at the detector (if a component of the displacement is along the dispersion axis of the spectrometer). A trapped particle moving between regions results in a sharp change in spectral output, with regions containing two traps resulting in overlapping spectra (i.e. cross-talk between readout from each trap). To reduce such cross-talk, the CCD regions for spectral readout can be reduced in size to match the vertical extent of the dispersed spectrum (typically 10 CCD tracks for a pinhole of 110*μ*m). Nevertheless, this example demonstrates the capability of the HOT-RMS instrument for truly simultaneous real-time HOT-RMS measurements of multiple trapped objects, even when simulatneously recording bright-field images with halogen lamp illumination.

**Figure 2:**
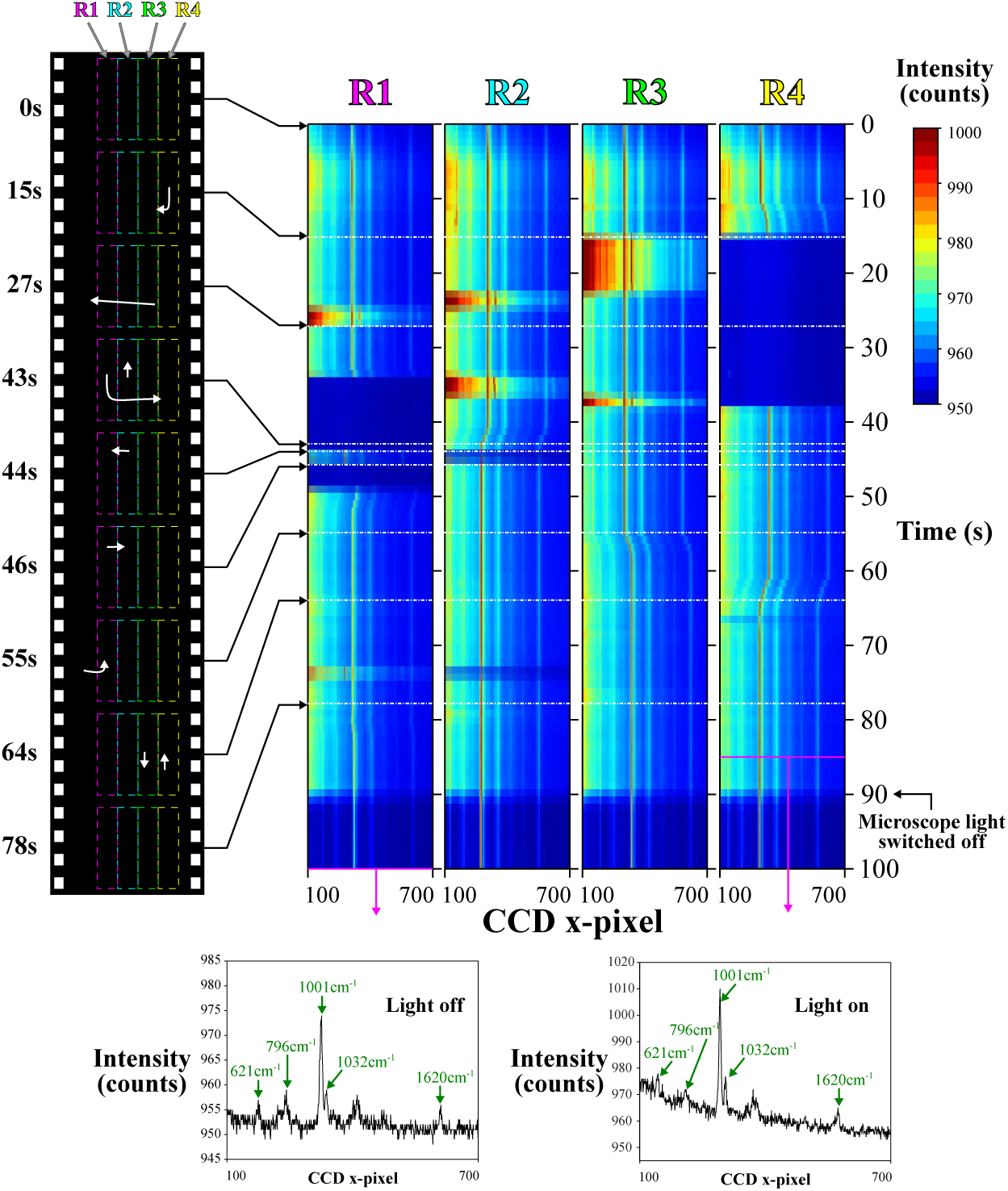
Demonstration of real-time interactive optical manipulation and RMS acquisition. The numbered regions marked by colored rectangles in the movie frames (from DMD camera) correspond to four CCD binning regions used to acquire Raman spectra (see Video 1). These frames show manipulation of four polystyrene beads (5μm) within the different regions. During optical manipulation, binned CCD acquisitions are obtained at 10 Hz (rate limited by mechanical shutter speed, exposure time 10ms), with each readout producing four Raman spectra, one from each region. The time-resolved spectra are shown in the four colormap images (color limits adjusted for clarity). For the first 90 seconds, measurements were carried out with the microscope halogen light illumination for bright-field imaging.

## HOT-RMS OF LIVING CELLS AND ORGANISMS

RMS measurements of live cells in solution typically involve using laser excitation powers around 20-200mW at the sample (at 785nm). As the spontaneous Raman effect is a relatively weak phenomenon and Raman scattering intensity scales linearly with excitation intensity, the excitation laser power would ideally be as large as possible. However above 200mW, many cells tend to be irreversibly damaged(1). This damage threshold is not universal, and can significantly vary depending on the exact optical properties of the cells under investigation, with mammalian erythrocytes (red blood cells: RBCs) for example being much more sensitive at 785nm excitation, as they exhibit an absorption resonance leading to a damage threshold at approximately 20mW(36). It is therefore essential to be able to control the laser power for each HOT-RMS laser trap such that excitation can be optimized whilst avoiding laser-induced damage and allowing quantitative comparison between measured Raman spectra. As only the total power (*P*_0_) of the laser spot array can be measured using a power meter (due to the microscopic separation of the spots), we estimate the power per trap (P_*i*_) from P_0_ using the equation

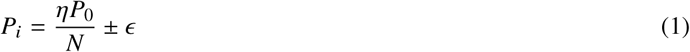

where *η* is the LC-SLM modulation efficiency, *N* is the total number of software-generated laser foci, and *ϵ* is an error term. In order to vary **P**_0_ itself, a variable attenuator was placed before the LC-SLM allowing *P*_0_ to span 0-800mW at the microscope objective. The *η* term results in an optical trap, which cannot be controlled in software, of power (1 - *η*)*P*_0_ fixed on the optical axis (for the LC-SLM model used here, *η* ≈ 0.9). The error term was estimated using two approaches, the first used non-saturated microscope camera images of different laser patterns focused at a quartz-air interface, and the second used Raman spectra from a uniform transparent polystyrene surface. The results from both approaches are presented in supplementary Figure S5, and result in an estimation for the relative error in *P_i_* of ≈ = 15%. One aspect overlooked by equation 1 is higher-order diffraction from the LC-SLM. This was minimized by adjusting the look-up table (LUT) in the RedTweezers software such that higher order diffraction was minimized (21).

Figure 3 shows examples of optically trapped cells with their correponding Raman spectra at different excitation/trapping powers. For RBCs which have a high absorption coefficient, resonant Raman spectra of hemezoin and hemoglobin are measured at 785nm excitation, enabling measurement of the multiple high SNR Raman spectra shown in Figure 3(a) at viable laser powers as estimated by equation 1 (36). For similarly sized cells such as yeast, excitation is possible at larger P_*i*_, so Raman spectra of similar SNR can be obtained with shorter acquisition times, as shown in Figure 3(b). It is also possible to measure smaller trapped cells such as bacteria at similar P_*i*_ to yeast as seen in Figure 3(c), though the integration time required is typically longer as bacteria have lower total biomass which will undergo Raman scattering within the laser focus.

**Figure 3:**
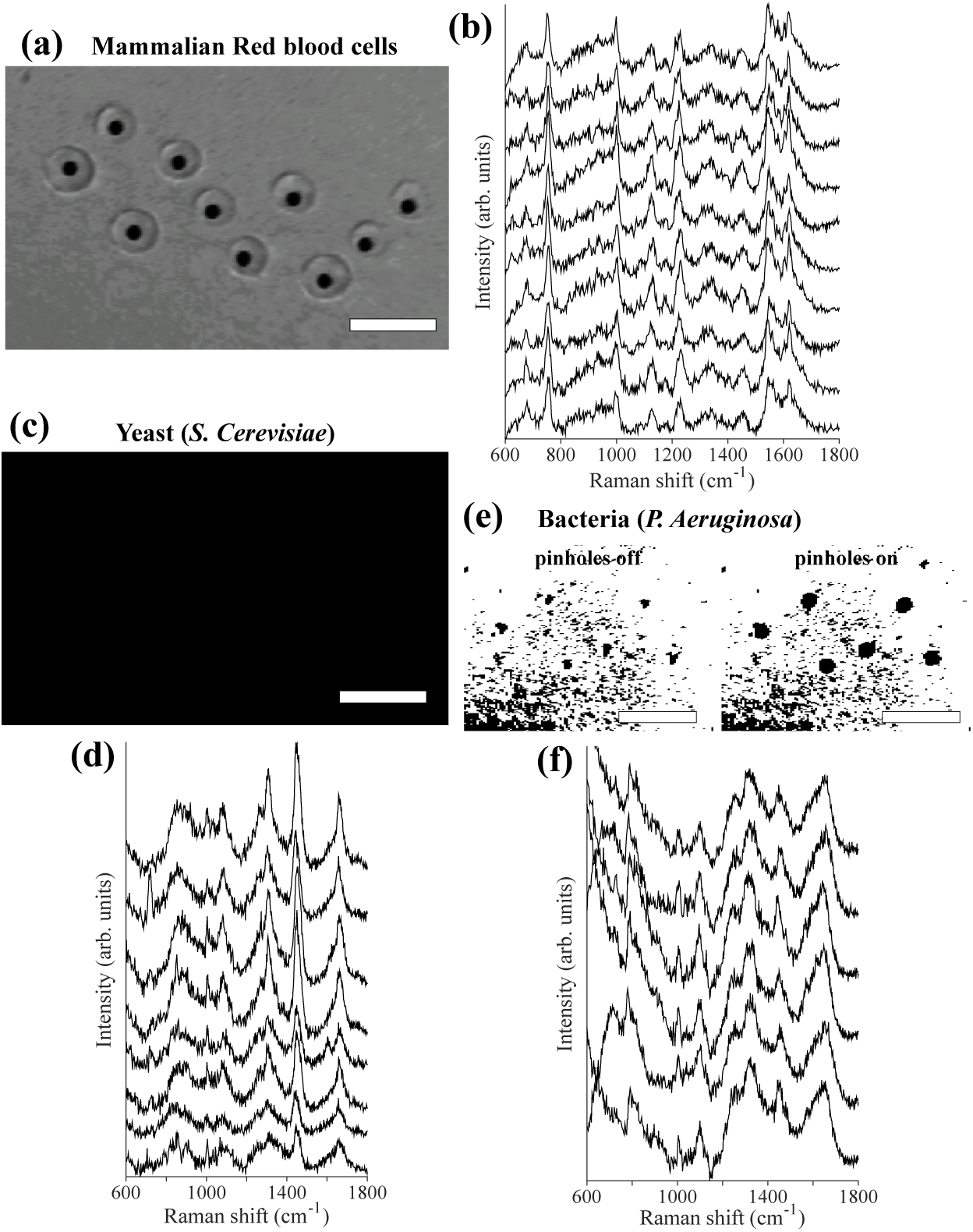
Examples of HOT-RMS multi-beam trapping measurements of various cell types, with DMD camera images (scale bars 10*μ*m) showing the fixed pinhole locations for measurement of the trapped cells (a,c,e) and Raman spectra from each trapping location (b,d,f; spectra vertically shifted for clarity). (a,b) Mammalian red blood cells (*Pi* = 3mW, 60s acquisition, 5th order polynomial baseline and quartz background subtraction). (c,d) *S. Cerevisiae* yeast (*P_i_* =50mW, 2s acquisition, 1st order polynomial baseline subtracted). (e,f) *P. Aeruginosa* bacteria (P_*i*_ =50mW, 10s acquisition, 1st order polynomial baseline subtraction)

Laser-damage mechanisms possible during RMS measurements may involve two-photon fluorescence, laser-induced heating, and photochemical reactions of particular light-sensitive compounds (e.g. porphyrins) within cells(1). For RMS at 785nm excitation, it is often the latter effect responsible for damage. In some cases, time-course Raman spectra of trapped cells may show spectral changes which indicate the mechanism of laser-induced damage. RBCs are one such case, as the resonant Raman signal of the hemoglobin provides sufficient SNR and specificity to monitor the laser-induced aggregation with a reasonable temporal resolution. Figure 4 shows RBCs used as a test case, whereby cells trapped with different values of *P*_*i*_, ranging from 3mW to 27mW, with time-course Raman spectra acquired starting from the moment the cells are captured in a trap. It can be seen in Figure 4(a) with *P*_*i*_ = 27mW (above the damage threshold) that the time-course spectra show a rapid spectral change over the course of 30s, with the Raman difference spectrum between the mean Raman spectrum from the first and last 5s of the time-course(4(b)). The differences show an increase in baseline indicating increased fluorescence, as well as a change in certain Raman band intensities, particularly the 1250cm^−1^band, which has previously been assigned as a denaturation and aggregation biomarker (36).

**Figure 4:**
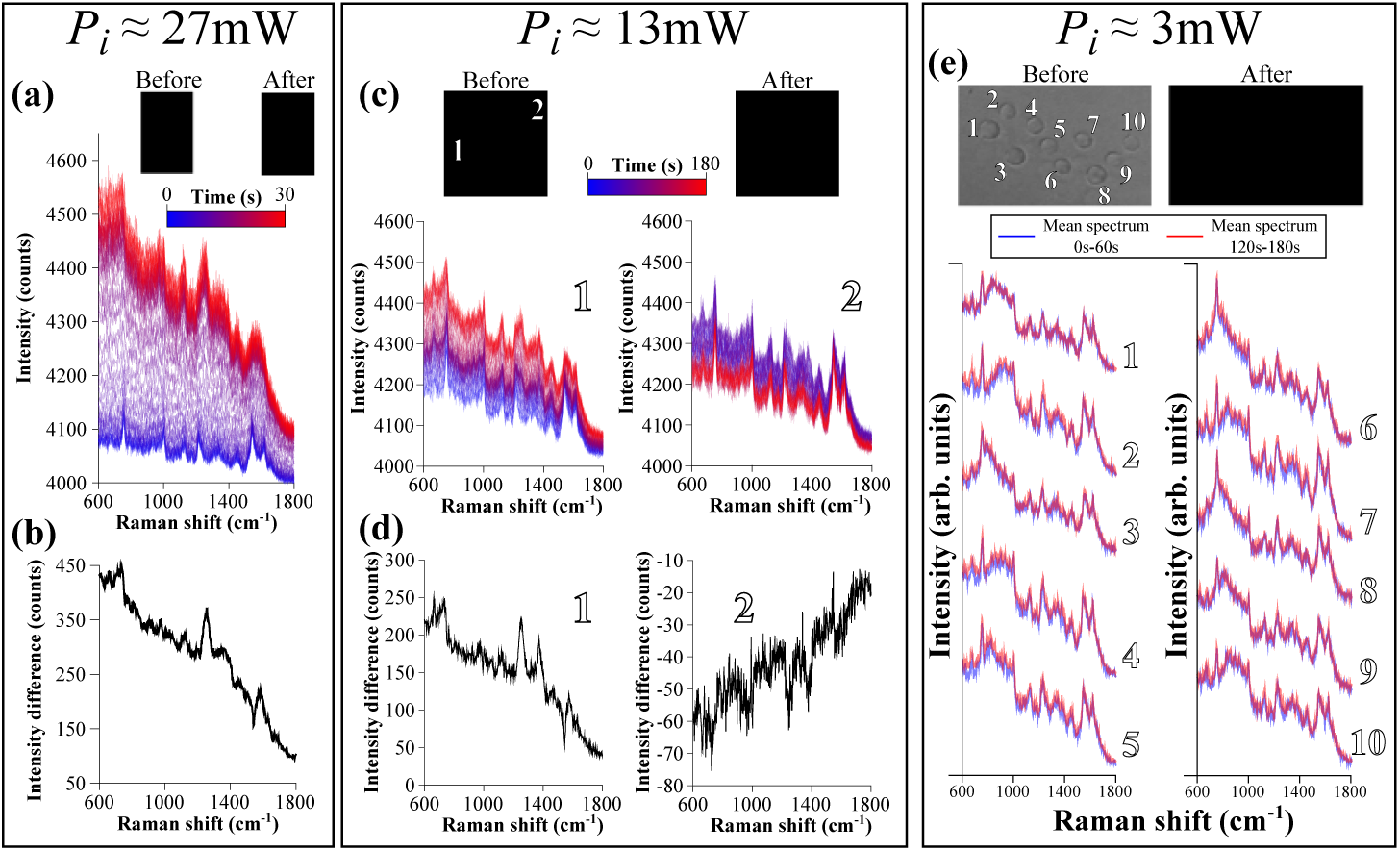
Control of laser power for HOT-Raman measurements using red blood cells (RBCs). (a) Time-course Raman spectra (raw data) for equine RBCs trapped at *P*_*i*_ =27mW (0.5s acquisition per spectrum). The Raman difference spectrum (mean of final 10 spectra minus mean of initial 10 Raman spectra) is also shown in (b). (c) Time-course Raman spectra for two RBCs trapped with *P*_*i*_ =13mW (2.5s acquisition per spectrum), with (d) showing Raman difference spectra from each cell (mean of final 10 minus mean of first 10 Raman spectra). (e) For ten RBCs trapped with P_*i*_ =3mW, the mean of the first and last 60s of a 180s time-course Raman spectra acquisition are shown.

Figure 4(c) shows a similar measurement with two cells trapped and measured simultaneously, with *P*_*i*_ = 13mW which is below the estimated laser damage threshold. The time-course Raman spectra over 180s show that cell 1 experiences similar denaturation and aggregation to that in Figure 4(a,b), albeit at a reduced rate, with the difference shown in Figure 4(d). In contrast, the Raman spectra from cell number 2 shows minimal change, with a slight decrease in signal, possibly due to a change in concentration of the RBC constituents due to heating or trapping forces. The difference between these two cells measured at the same expected value of *P*_*i*_ could be due to the error ≈ =15% mentioned in equation 1 leading to one beam being above the damage threshold with the other below the threshold, or possibly from inter-cellular differences such as the cell size or a different concentration of hemoglobin within the cell.

Figure 4(e) shows a measurement with *P*_*i*_ =3mW for ten simultaneously trapped RBCs. The spectra shown are averages of the first and last 60s of the time-course Raman spectra (over 180s total), which show no measurable differences above the noise level of the measurement, suggesting immediate laser-induced damage has been avoided for all ten cells in the trap.

The measurements in Figures 3 and 3 demonstrate that *P*_*i*_ can be sufficiently controlled to optimize measurements for cells on a case-by-case basis, and that optical throughput is sufficient to allow time-course monitoring of laser-induced damage in the case of RBCs. While most other cell types do not exhibit such a strong resonant Raman response, they can be expected to be more resilient to laser exposure at higher powers assuming their absorption cross-section is lower. One example of this is yeast cells, which exhibit a useful Raman band at 1602cm^−1^which has been used as a live/dead biomarker linked to metabolic activity(5). Using a similar approach to that shown in Figure 4, the HOT-RMS instrument was used to estimate the laser-power damage transition using this 1602cm^−1^ biomarker for yeast (see supplementary Figure S6).

## HOT-RMS FOR INDUCING AND MEASURING IMMUNOLOGICAL SYNAPSES

To demonstrate the unique capability of the HOT-RMS instrument, we manipulated multiple live T cells, bringing them controllably into contact with a dendritic cell (DC) to initiate the formation of an immunological synapse (IS) junction, and measuring space- and time-resolved Raman spectra.

Figure 5 shows an example of an induced interaction between multiple T cells and DCs using HOT with measurement at different locations and time points using RMS. All HOT and RMS measurements were acquired with 6 active traps at any one time (*P*_*i*_ ≈50mW), and the laser was blocked from the sample between trapping/Raman acquisitions. A time-course Raman acquisition for a trapped T cell in these conditions is shown in supplementary Figure S7, with no significant laser-induced damage effects observed. The example interaction shown in Figure 5 involves a DC with an existing connection to a T cell (labelled TC1). The microscope image shown in Figure 5 (a) shows the process of initiating a second interaction between the DC and an optically trapped T cell (TC2) at time t *=* 0. Attachment was verified by attempting to trap and pull off the T cells after waiting for a short time. A cell was considered attached once the laser trap was incapable of moving the T cell. Once TC2 was attached, Raman spectra from various points were acquired over the course of 5 minutes, with selected spectra shown from the labelled positions in the schematic in Figure 5 (a), focusing particularly on T cells (green), DC edges (purple) and junctions (blue). It can be seen that the T cell spectra (TC1 and TC2) contain relatively strong Raman contributions from nucleic acids (788cm^−1^and 1090cm^−1^) and proteins (1004cm^−1^and 1660cm^−1^), in agreement with previous studies(9,10). In contrast, DCs generally have a weaker Raman signal which varies spatially, with certain regions showing lipid droplet signals (E1,E3). Figure 5 (b) shows the interacting system after t = +1 hour, when two additional T cells were trapped and brought towards the DC for a further interaction. After this, several Raman spectra were obtained in a similar manner to Figure 5(a) at the locations shown in the schematic in Figure 5(b). Similar observations can be made to the measurements shortly after t *=* 0, with the exception of the newly formed junctions, J3 and J4, between the DC and the T cells TC3 and TC4.

**Figure 5:**
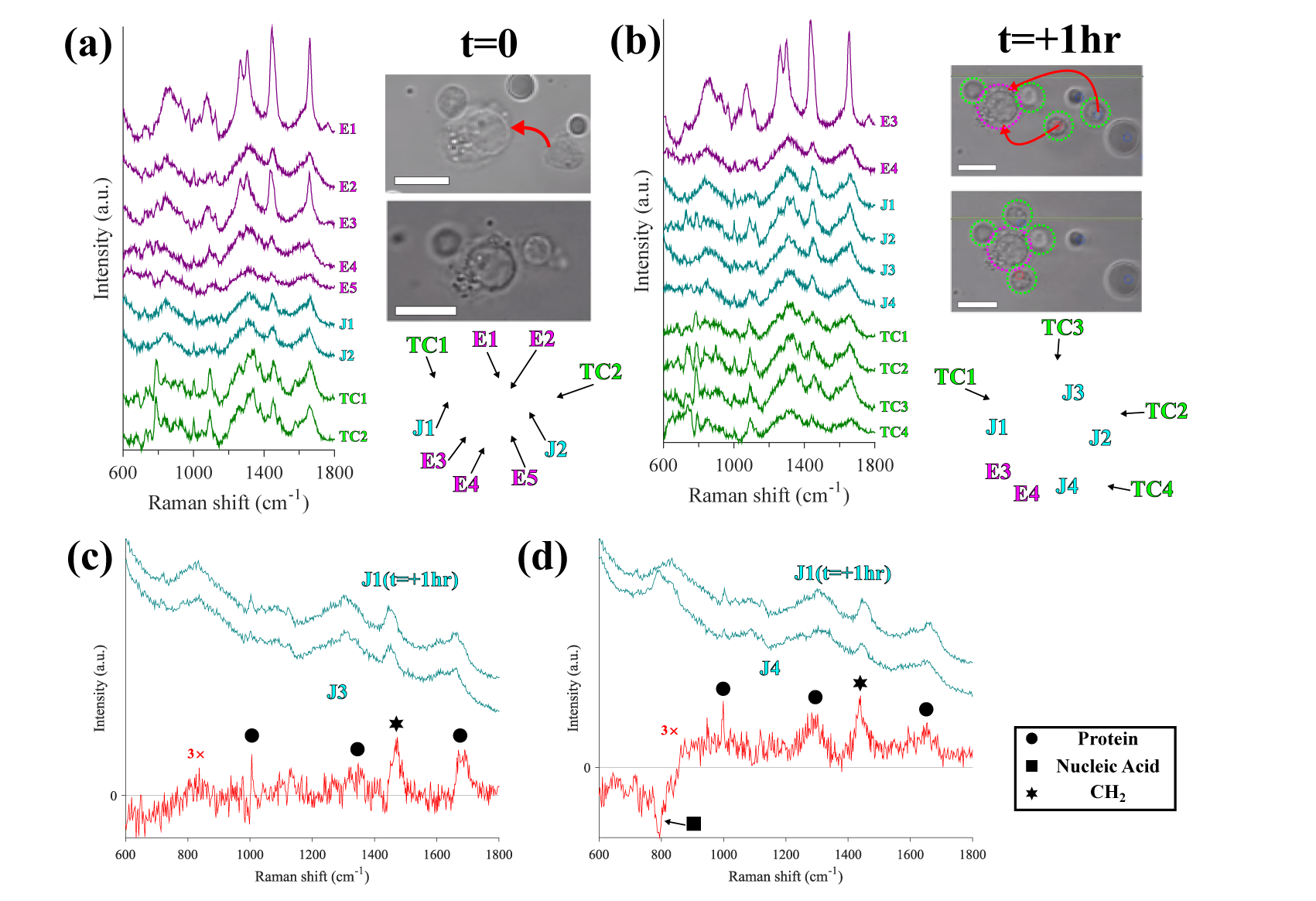
Formation of an immunological synapse (IS) between an adherent dendritic cell (DC) and optically trapped T cells monitored by RMS. (a) Raman spectra and microscope images at t = 0 when a DC with one T cell already interacting has another T cell brought into contact using the HOT-Raman instrument. The schematic shows the locations from which Raman spectra were acquired (20s acquisition). (b) Raman spectra and microscope images showing further induced interactions with two more trapped T cells brought into contact with the same DC in (a) at t = +1hr, with a second schematic showing estimated sampling locations. (c) and (d) show Raman difference spectra between the oldest IS junction (J1) and the new junctions J3 and J4 (measured at t=+1hr shortly after their formation). Raman spectra in (a) and (b) have a linear baseline and quartz background subtracted, wheras (c) and (d) show raw Raman spectral data. All scale bars 10μm.

The Raman spectra from the various locations shown in Figure 5(a) and (b) can be compared to identify dynamic changes occuring in the interacting cell system. The Raman bands from the T cells and DC edges appear to be consistent at both timepoints. However the differences between the Raman spectra measured at the junctions suggest molecular differences, in particular protein content. Figure 5 (c) and (d) show the Raman difference spectra between J1 (the earliest junction formed between a T cell and the DC, as measured at t = 0) and J3 and J4 respectively induced by HOT at time t = + 1hr. The difference appears to mainly contain Raman bands assigned to proteins, particularly the prominent 1004cm^−1^band (breathing mode of phenylalanine ring) which appears larger relative to the 1450cm^−1^(CH_2_ and CH_3_ deformation vibrations) in the difference spectrum compared to the individual Raman spectra, suggesting the junctions which formed earliest have a larger protein content. Previous studies of fixed DCs and T cells comparing Raman spectral imaging and fluorescence staining of actin indicated that such changes in protein level may be linked to actin polarisation at the IS (10). These differences may also be due to an increased measurement volume between the sampling locations, and would require thickness/morphology measurement to conclusively determine which case these measurements fall into(14).

This demonstration of a HOT-RMS measurement between multiple dynamically interacting cells demonstrates the capability of the instrument for label-free molecular sensing. As optical trapping can be used for T cell manipulation, interactions between the immune cells can be monitored from the start, with endogenous molecular changes at the formed junctions detected by RMS. Such capability could offer previously unattainable insight into the kinetics of biochemical changes during all time-points of cell-cell interactions.

## HOT-RMS FOR CONTROLLED PARASITE INFECTION AND SPECTROSCOPIC MEASUREMENT

While RMS is often utilized to measure endogenous biomolecules, substitutions with a stable isotope such as deuterium (D) or ^13^C, which will modify the frequency of a vibrational mode, can be detected in a Raman spectrum. These labels can be used to show uptake and metabolism in cells(37) or the transfer of amino acids between a host and pathogen(11). Here, we demonstrate a HOT-RMS experiment of a stable-isotope labelled cell system. The interaction between human brain microvascular endothelial cells (hBMECs) cultured in media with deuterated phenylalanine and *T. gondii* parasites containing normal phenylalanine can be initiated by HOT, and monitored by RMS. Previous RMS studies have involved observations of this infection with the earliest times starting at several hours, with the initial stages of infection not observed as infection was left to progress with no control(11). As we are able to initiate the interaction by optically trapping *T. Gondii,* the infection can be monitored from the time of initial contact, with multiple Raman spectra measurable from different points of interest simultaneously. Figure 6 shows an example measurement of the hBMEC-*T. Gondii* interaction on the HOT-RMS instrument. The video frames shown in Figure 6(a) illustrate the manipulation of *T. Gondii* parasites towards an adherent hBMEC on the quartz coverslip surface. Multiple isolated *T. Gondii* were trapped and brought into contact with a hBMEC cell membrane surface, with typically fewer than 10% appearing to attach after several minutes of observation. Time-course measurements of several *T. Gondii* showed no measurable adverse effects of trapping on the parasites (see supplementary Figure S8). Once cells are brought into contact, the laser is blocked from the sample until RMS measurement. In a succesful attachment, the apical end of the *T. Gondii* appears fixed at the cell membrane surface of the hBMEC close to the position it was manipulated towards at t=0 using the optical traps, as can be seen in the frame at t=+15min. At this point, the parasite appears able to be partially trapped, whilst tethered at one end to the hBMEC. At t=+85min, the parasite has entered the cytoplasm of the hBMEC and is no longer able to be moved by the optical trap. Internalized parasites were distinguished from those simply above/below the cell from their absence of Brownian motion visible on the microscope camera.

**Figure 6:**
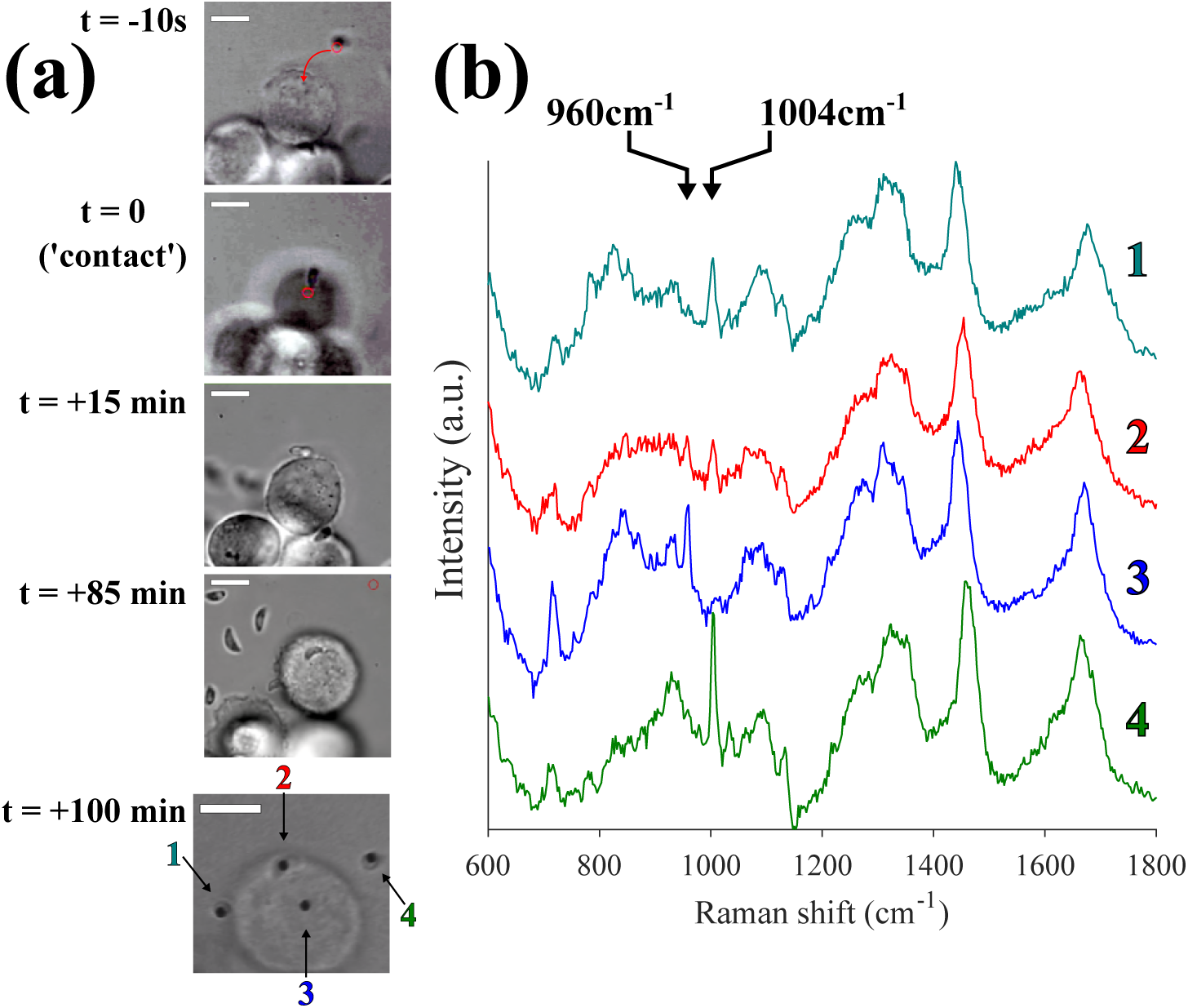
HOT-Raman instrument to initiate and measure Raman spectra of a single *T. Gondii* infection in hBMEC cells. (a) Microscope camera frames showing an optically trapped T-gondii brought towards a hBMEC cell. The last frame shows a DMD camera image of four measurement points corresponding to (1) *T. Gondii* attached to hBMEC surface, (2) *T. Gondii* at the early stage of infection in hBMEC, (3) hBMEC cytoplasm and (4) isolated trapped *T. Gondii.* The Raman spectra corresponding to the points (1)-(4) are shown in (b) (baseline subtracted raw data, spectra shifted for clarity, acquisition time 1 minute, *P_i_ =* 30mW). All scale bars in (a) are 10μm.

At t=100min, Raman spectra were acquired from four different locations within the host-pathogen system simultaneously *(*P*_i_ =* 30mW, 6 beams present at sample, two unused), as can be observed in Figure 6(b). It can be seen that the isolated parasite (4) and the attached parasite at the early stage of infection (1) exhibit the standard 1004cm^−1^ phenyl Raman band, with no observable deuterated phenylalanine peak at 960cm^−1^ in either spectrum. In contrast, the Raman spectrum from the cytoplasm of the hBMEC (3) shows only the presence of the deuterated phenylalanine Raman band at 960cm^−1^ as expected. The internalized *T. Gondii* parasite (2) appears to contain contributions from both 960cm^−1^and 1004cm^−1^Raman bands. This may indicate that the exchange has begun within the 100 minutes, or that the laser is exciting Raman scattering from a volume containing both non-labelled phenylalanine from the parasite, and labelled proteins from the host cytoplasm. Therefore a full calibration of the instrument should be performed prior to attempting a quantitative study of such a system.

This experiment demonstrates that in addition to label-free RMS, labelled methods with increased specificity can also be utilized in an interacting cell system, which could track the transport of molecules between cells.

## CONCLUSION

We have demonstrated that HOT-RMS is a promising tool for simultaneous non-invasive manipulation and RMS molecular measurement of interacting living organisms. Raman spectra with spectral resolution from diffraction limited points were able to be obtained from multiple trapped objects simultaneously in arbitrary locations in the field of view, with high temporal resolution. We have demonstrated the general applicability of such an instrument, from measuring yeast, blood and mammalian immune cells, to pathogens such as bacteria and intracellular parasites. The capability for high-resolution time-course RMS spectra and accurate laser trap power control can also be used to assess laser-induced damage thresholds and measurement viability. We finally presented examples of interacting cell systems; with DC and T cells forming immunological synapses, and a *T. Gondii* parasite with an amino-acid isotope-labelled hBMEC host cell.

These results are a unique capability of the HOT-Raman instrument, which allows rapid, flexible, multiplexed measurement and manipulation of live cell systems, enabled by the LC-SLM and DMD technologies.

## AUTHOR CONTRIBUTIONS

FS, GG, MP and IN designed the instrument. FS built instrument, carried out experiments, analyzed data, and prepared all figures. HME provided materials for and assisted with host-pathogen experiments, which were designed by FS, HME and IN. AMG and DA provided materials and assisted with immune cell experiments, which were designed by FS, DA, AMG and IN. FS, HME, GG, AMG and IN wrote the paper.

## ACKNOWLEDGMENTS

The authors would like to thank Professor Paul Williams and Dr. James Brown for providing samples of planktonic *P. Aeruginosa* bacteria for trapping measurements. F.S. acknowledges the University of Nottingham for the award of a UK Engineering and Physical Sciences Research Council (EPSRC) doctoral prize (Grant number: EP/M506588/1). I.N. holds an established career fellowship from EPSRC (Grant number: EP/L025620/1).

## SUPPLEMENTARY MATERIAL

An online supplement to this article can be found by visiting BJ Online at http://www.biophysj.org.

